# Evolution of a novel female reproductive strategy in *Drosophila melanogaster* populations subjected to long term protein restriction

**DOI:** 10.1101/2021.10.06.463438

**Authors:** Purbasha Dasgupta, Subhasish Halder, Debapriya Dari, P. Nabeel, Sai Samhitha Vajja, Bodhisatta Nandy

**Author notes:** Corresponding author, contact number. **Author contributions:** BN conceived the idea, designed the experiment, analysed and interpreted the data, wrote the manuscript. PD designed and performed the experiments, analysed data, prepared the manuscript. SH, DD, NP, and SSV performed various assays and collected data. BN and PD revised the manuscript and wrote the responses to reviewers’ comments.

## Abstract

Reproductive output is often constrained by availability of macronutrients, especially protein. Long term protein restriction, therefore, is expected to select for traits that maximize reproductive output in spite of such nutritional challenge. We subjected four replicate populations of *Drosophila melanogaster* to a complete deprivation of live-yeast supplement, thereby mimicking a protein restricted ecology. As yeast supplement is a key source of protein, such an ecology is expected to strongly limit reproductive output, especially in females. Following 24 generations of experimental evolution, compared to their matched controls, females from experimentally evolved populations showed increase in reproductive output early in life, both in presence and absence of yeast supplement. The observed increase in reproductive output was not associated with any accommodating alteration in egg size; and development time, pre-adult survivorship, and body mass at eclosion of the progeny. There was no evidence for evolution of lifespan and lifelong cumulative reproductive output in females. However, females from experiment regime were found to have a significantly faster rate of reproductive senescence, as indicated by a faster rate of age related decline in reproductive output following the attainment of the reproductive peak. Therefore, adaptation to yeast deprivation ecology in our study involved a novel reproductive strategy whereby females attained higher reproductive output early in life followed by faster reproductive aging. To the best of our knowledge, this set of results is one of the most clear demonstration of optimization of fitness by fine tuning of reproductive schedule during adaptation to a prolonged nutritional deprivation.

**Supplementary information:** A separate file that includes three figures and six tables.

## 1. Introduction

The ability of an organism to reproduce in spite of nutritional challenges is essential to adaptation in a given ecology (Kolss et al 2009, Zhang et al. 2019). Beyond providing the essential energy to sustain life, nutrients, especially macronutrients such as proteins and carbohydrates, determine survival by affecting resistance to stress and other ecological challenges, including pathogens (Chippindale 1993, Field et al. 2002, Andersen et al. 2010, Sisodia and Singh 2012, Miller and Cotter 2017, Weldon et al. 2019). Access to macronutrients also affect reproductive output and is central to a variety of trade-offs concerning reproduction (Chippindale 1993, Tatar & Carey 1995, Chapman & Partridge 1996, Marden et al. 2003, Adler & Bonduriansky 2014). This is evident in response to a form of nutritional challenge, wherein one or more essential macronutrients is restricted without causing malnutrition, organisms have been found to elicit a complex physiological response that culminates in the partial or complete shutdown of reproduction, resulting in an extension of lifespan (Mair et al. 2005, Partridge et al. 2005, Nakagawa et al. 2012, Fontana & Partridge 2015). Temporarily shutting down reproduction could be advantageous if it allows the organism to tide over the period of nutritional shortage, following which reproduction can resume. However, such a reproductive strategy is expected to be detrimental if nutritional shortage is long lasting, spanning generations. There is no reason to not expect such long term nutritional shortage. It is not clear how organisms adaptively optimize reproduction when faced with such selective pressure (Adler & Bonduriansky 2014). Perhaps the most famous example of adaptation to such dietary shortage is the evolution of Finch populations in the Galapagos archipelago following an intense selection imposed by a prolonged period of draught that led to a severe qualitative and quantitative depletion of the food supply, resulting in strong natural selection (Boag & Grant 1981). It resulted in a complex evolutionary response in morphology, physiology and behaviour of the finch populations (for example, see Schluter & Grant 1984). Thus, undoubtedly diet can drive adaptive evolution. Recent studies on nutritional deprivation have shown that deprivation of specific macronutrients, especially protein, can pose specific effects on fitness components – limiting reproduction (Maklakov et al. 2008, Lee et al. 2008, Fanson et al. 2009, Fanson and Taylor 2012, Short et al. 2020), stress resistance (Sisodia & Singh 2012), memory and learning (Mery & Kawecki 2005). However, surprisingly little is known about how populations evolving under nutritional deprivation, specifically protein deprivation, optimize reproduction. In addition, there is also a serious dearth of literature on genetic variance and covariance for traits that could allow or impede such adaptation (but see Rikke et al. 2010, Jin et al. 2020).

Apart from other model and non-model organisms, *Drosophila melanogaster* has been extensively used to investigate the plasticity in life history and the underlying physiological response to dietary restriction. Among the macronutrients, while carbohydrate is the primary caloric source, protein is a major determinant of reproductive output. Both in laboratory cultures and in wild populations, Drosophilid fruit flies acquire protein by foraging on yeast (Begon 1982, Starmer 1982, Chippindale et al., 1993; Anagnostou, Dorsch & Rohlfs 2010). Restricted access to yeast, therefore, lead to protein starvation, and is known to result in reduced reproductive output, especially in females (Lee et al. 2008). Yeast deprivation in laboratory systems of *Drosophila* sp. is usually achieved either by limiting yeast supplement on food (Partridge et al. 1987, Chippindale et al.1993) or by reducing the amount of yeast in the food (Chapman and Partridge, 1996) – both leading to reduced fecundity in females. Such reduction in reproduction, triggered by dietary restriction, has been frequently found to be accompanied with increased lifespan, inspiring a large body of experimental research. There is, now, ample evidence to suggest that such lifespan extension is almost entirely due to yeast restriction and not caloric or carbohydrate restriction (Zimmerman et al. 2003, Mair et al. 2005, Piper et al. 2005). Further, alteration in the dietary yeast to sucrose ratio, within certain limits, without changing the caloric value of food has been found to generate a physiological shift away from reproduction (Lee et al. 2008). Other nutritional stresses, including extreme starvation, have also been shown to lead to a physiological shift resulting in extension of lifespan, often at the cost of reduced fecundity in females (Hoffmann & Harshman 1999, Parkash & Munjal 2000, Partridge et al. 2005, Rion & Kawecki 2007). Although adaptive during a transient nutritional deprivation (Shanley & Kirkwood 1999, Adler & Bonduriansky 2014, Moatt et al. 2020), such a physiological response of diversion of resources away from reproduction is an unlikely solution for prolonged nutritional scarcity. In such case, natural selection may favour a higher investment in reproduction even at the cost of survival, especially for short lived organisms (Williams 1957, Bonduriansky et al. 2008).

Experimental evolution using fruit flies has been a powerful approach to study a variety of evolutionary genetic problems, including adaptation to nutritional challenges (Prasad and Joshi 2003, Garland and Rose 2009). Kolss et al. (2009) showed that *D. melanogaster* populations can adapt to nutritional deprivation by evolving better egg-adult survivorship and faster development. More recent studies investigated adaptation to yeast-restricted adult diet to find that males under such a regime evolve higher reproductive output on a DR food without any significant change in lifespan (Zajitschek, 2016). However, the females in the same study on the other hand, evolved shorter lifespan with no alteration in their reproductive output (Zajitschek, 2019). Importantly, the outcome of experimental evolution critically depends on the specific context of the experimental selection pressure and the definition of fitness in the experimental population (Rose et al. 1996, Garland and Rose 2009). Hence, the issue of reproductive adaptation under nutritional deprivation is only beginning to emerge as a research programme with plenty of open questions.

Here we investigated adaptation of *D. melanogaster* populations to yeast deprivation specifically targeting yeast supplement in their diet instead of heat-killed yeast in their food. Importantly, besides affecting female reproductive output, yeast supplement is known to impact adult lifespan and starvation resistance (Chippindale et al. 1993), oviposition site preference (Becher et al 2012), and may even affect offspring survival and fitness components (Rohlfs & Hoffmeister 2005, Anagnostou, Dorsch & Rohlfs 2010). Given the strong fitness consequence of access to yeast supplement, the lack of experimental data on adaptation to yeast deprivation in the available literature is rather surprising. Interestingly, because yeast acts as dietary protein supplement primarily for females (Chippindale et al. 1993, Nandy et al. 2012), it is possible for yeast deprivation to represent a sex-specific selective environment, which can lead to sex specific adaptations as found by Zajitschek et al. (2016, 2019).

To investigate adaptation to yeast deprivation, we subjected four replicate populations of *D. melanogaster* to a complete withdrawal of yeast supplement from their diet. After 23 generations of experimental evolution, we investigated the effect of experimental evolution on (a) female fecundity in absence of yeast supplement as well as under varying access to the same, (b) a measure of female fitness, (c) pre-adult traits that can affect female fecundity, and (d) longevity and age specific reproductive output in females. Keeping the adaptation theory in mind, we predicted female fecundity and reproductive output to improve in the selected populations compared to the ancestral base populations, triggering correlated changes in the life history.

## 2. Materials and methods

### Study populations and selection regime

Four new experimental populations (referred to as YLB_1-4_ here on) were derived from four replicate baseline populations BL_1-4_ (Figure S1). The BL populations (Nandy et al. 2016) are outbred, wild type populations of *D. melanogaster*. They are maintained under a 14-day discrete generation cycle, under 24-hour light, at 25 °C (±1) ambient temperature, on a standard banana-sugarcane jaggery-yeast medium, with a population size of ∼2,800. Food is prepared by boiling mashed ripe banana, barley, sugarcane jaggery, and baker’s yeast in water. Agar-agar is used as solidifying agent and p-hydroxybenzoic acid dissolved in ethanol is used as a preservative. The complete recipe with the cooking method can be found in Supplementary information (Table S1). Flies are grown in standard vials (25 mm diameter × 96 mm height) for 12 days. Larval density is controlled at ∼70 per vial with 8 ml food and 40 such vials are maintained for each replicate population. On day 12 of the generation cycle,

∼2,800 adults are transferred to each plexiglass cage (23cm × 20cm × 15 cm) with a food plate smeared with an *ad-lib* quantity of live yeast (a paste made with water). On day 14 post-egg collection, the flies are given fresh food for oviposition and eggs laid during an 18-hour window (fitness window) are collected to start the next generation. BL populations were maintained in the laboratory under this regime for almost five years prior to the derivation of the experimental YLB populations. From each of the four replicate populations of BL (BL_1-4_), a corresponding YLB population (YLB_1-4_) was derived by collecting 40 vials of eggs with a standard egg density (i.e., approximately 2800 eggs were used to found each YLB population). The maintenance of the YLB populations is identical to that of the BL except for the absence of live-yeast supplement in the YLB ecology. All assays reported here were conducted after 23-25 generations of experimental evolution. To equalize any potential non-genetic parental effect, all populations were passed through one generation of common rearing ecology (Rose 1984). During this, a subset of the experimental populations was generated by collecting eggs from the stock BL_x_ and YLB_x_ populations. These populations were maintained under a regime that exactly matched that of the BL-regime (i.e. both BL and YLB were provided with *ad-lib* live-yeast). Experimental flies were generated by collecting eggs from these subset populations. During the assays, all fly handling, including the collection of virgin flies, wherever necessary, was done under light CO_2_ anaesthesia. A schematic representation of the overall experimental plan can be found in the supplementary information (Fig. S2).

### Experiment 1: Juvenile traits and early-life fecundity in females

After 23 generations of experimental evolution, we assessed whether the YLB populations have adaptively diverged from the ancestral BL populations. Our initial observation suggested fecundity to be the most likely target trait of selection (data not shown). Therefore, as the first set of assays, we quantified female early-life fecundity under a range of yeast concentrations. In addition, since fecundity is often a function of body size and related juvenile traits, we measured these traits.

Female early-life fecundity and three juvenile traits – pre-adult (egg-to-adult) development time, juvenile (egg-to-adult) survivorship and dry weight at eclosion were measured between 23-25 generations. Following one generation of common rearing (see above), eggs were collected from the population cages in exact numbers. For egg-collection, two small food plates were introduced in a cage, and were left undisturbed for about one hour to allow oviposition. Such a short window of oviposition ensured the developmental synchrony of the eggs collected for the experiment. Eggs were randomly scooped out from these oviposition plates, and transferred to an egg-collection surface made of solidified agar gel (1% Agar-agar solution) using a fine brush. Eggs were carefully counted and a small piece of Agar gel containing 70 eggs was cut out and transferred to a fresh food vial having 8ml food. Ten such vials were set up for each population. Each vial was labelled individually and placed into the incubator. As an added precaution, the vials in the incubator were shuffled twice every day to randomize any positional effect. The vials were monitored carefully for the signs of eclosion. Beginning from the eclosion of the first individual, the eclosing adults were counted and sexed every 6 hours until all the pupae eclosed. This data was used to derive the average measure of egg-to-adult development time (i.e., pre-adult development time) for males and females in each vial. Out of the 70 eggs initially cultured, the proportion that finally eclosed from a vial was considered as the measure of egg-to-adult survivorship (i.e., juvenile survivorship). Three vials (one from the YLB_1_; two from BL_2_) were excluded from the analysis due to mishandling. For each population (BL_x_ and YLB_x_), 50 males and 50 females were collected for dry whole body weight measurement. They were collected within 6-hours of eclosion, and were immediately frozen at -20 °C. They were later dried at 60 °C for 48 hr and weighed in groups of five using Shimadzu AUW220D to the nearest 0.01 mg. The mean body weight of each group was calculated and taken as a unit of analysis.

For early-life fecundity assay, flies were raised and collected as virgins as mentioned above. Virgin flies were held in same-sex vials in groups of 12 in each vial. On the day12 post-egg collection (i.e., corresponding to 2-3 day post-eclosion), one vial each of males and females (i.e., 12 males and 12 females) belonging to the same regime, were transferred to a fresh food vial assigned to one of the three live-yeast supplementation treatments -No yeast (N), Limiting yeast (L: each vial containing 4.5 mg yeast suspended in 50 μl of water) and High yeast (H: each vial containing 9 mg yeast suspended in 50 μl of water). Twelve such vials were set up per yeast supplementation treatment for a population. For setting up L and H treatments, yeast was dissolved in tap water to make a suspension. It was then dispensed at a point on the rim of the food meniscus in a vial to ensure unfettered access to both food and the live-yeast. The suspension was allowed to air-dry before the introduction of the experimental flies. The males and the females were allowed to interact in these vials for 48 hours. Following this, on day 4-5 post-eclosion, flies in each treatment vial were anaesthetized under light CO_2_ -six females were randomly separated and transferred to a vial for egg size measurement, while the other six were individually transferred to six oviposition tubes (12 × 75 mm) for fecundity measurement. For the former, females were allowed to oviposit for 16 hours in a vial, following which they were discarded. The vial with eggs was frozen at -20 °C for a minimum of one day. For egg size measurement, eggs were mounted on a glass slide on their dorsal side and photographed using a stereozoom microscope and the area of the two-dimensional elliptical outline of the eggs were measured in Image J software. This area was taken as a proxy for the size of each egg. A given egg was measured three times and the average of these three measurements was taken as the unit of analysis. Females were allowed to oviposit in these tubes for 16 hours. Females from the oviposition tubes were then discarded and number of eggs was counted. We removed a few vials from the analysis due to a variety of handling errors, including escaped flies and broken tube/vial, during the assay. The final number of vials used for the analysis ranged from 9-12 vials per live-yeast treatment for a population.

#### Experiment 2: Female × male effect on female fitness

To further assess the response of female reproductive output, and eventually fitness, we quantified female early-life reproductive output (progeny count = number of surviving adult offspring produced by a female) during a 16-hour window on the day 4-5 post-eclosion. Given that most of the flies eclose during day9-10 (i.e., 9-10^th^ day of the generation cycle starting from egg stage), this window corresponded to the 18-hour fitness window on the 14^th^ day of the generation cycle and therefore, is a good measure of Darwinian fitness in this system (see Rice et al. 2006 for the measure of fitness in laboratory adapted populations). Importantly, female fitness, especially in *D. melanogaster*, is affected by both female fecundity and male factors such as seminal fluid composition (Chapman et al. 1995; Arnqvist & Nilsson 2000; Wolfner 2009). Therefore, it is important to assess male and female specific effect of selection on female fitness. To test this effect, we combined YLB and BL males and females in a full factorial design and measured early-life reproductive output (i.e., progeny count) of females in the above-mentioned protocol. These assays were carried out after 26-29 generations of selection. Experimental flies were generated following a protocol similar to that followed in Experiment 1. Males and females were collected as virgins (on around day 9-10 post-egg collection) and held in single-sex vials in groups of six in each vial until day 12 post-egg collection. On the day12 of the generation cycle (i.e., 2-3 days post-eclosion), sexes were combined to set up adult interaction vials having six individuals of each sex in a vial. Four types of combinations (viz., female×male: BL×YLB; BL×BL; YLB×YLB; YLB×BL), each having 15 replicate vials, were set up. A few flies escaped during the setting up of one vial in the third block with YLB_3_×YLB_3_ combination and hence, it was removed from the experiment. These adult interaction vials were left undisturbed for 48 hours. On the day 14, females from each of these vials were sorted under light CO_2_anaesthesia, and individually transferred to oviposition test tubes (one female per tube) where they were allowed to oviposit for 17 hours. After 17 hours, females from the oviposition tubes were discarded and the tubes were kept in the incubator for the development and emergence of the progeny. After the emergence of all the progeny, the test tubes were frozen at -20 °C for the final count.

#### Experiment 3: Longevity and age-specific fecundity

To test reproduction vs. lifespan trade-off theory, longevity and fecundity of the experimental females were measured after 26 generations of selection. To assess age related changes in fecundity, we measured fecundity of the test females in regular intervals. Similar to the other assays described above, all populations were passed through one generation of common rearing. The eggs were collected from the common rearing cages and cultured under standard density of ∼70 per 6-8ml food in a vial. Adult flies were collected as virgins within six hours of eclosion and held in single-sex vials in groups of 12 in standard food vials. These vials were left undisturbed for two days. On the day 3 post eclosion (i.e., on the day 12 post egg collection), experimental vials were set up by transferring the flies into fresh food vials. Two sets of vials were set up for each population – (a) Continuously exposed (CE): 12 males and 12 females placed (without anaesthesia) in a vial and (b) Virgin females (VF):12 virgin females placed in a vial. For a population, 12 vials were set up for each of CE and VF. Except for the fecundity count days (see below), flies were transferred into fresh food vials every alternate day without anaesthesia. Dead flies were sexed and counted during every transfer. In the course of the repeated vial-to-vial transfers, a few flies escaped at various time points. These flies were right censored for the survivorship analysis. For analysis of mean longevity, vials where more than four flies escaped were removed from the final analysis. Thus mean longevity analysis included data from 8-12 vials per treatment for each population. In this study, we have focused on the response to selection on female longevity only.

Fecundity of the CE females were measured at regular intervals. Starting from day5 post-eclosion (i.e. day 14 post-egg collection), every fifth day, the flies were transferred into fresh food vials (fecundity vials), where they were allowed a window of 17-hours before being transferred to fresh food vials again. The number of eggs in these fecundity vials was counted. This was done at day5, 10, 15, 20, 25, 30, 35, 40, 45, 50, 55, and 60 (by this time-point all females had died).For each time point, the number of eggs in a vial was divided by the number of surviving females to calculate per female fecundity. This was taken as the unit of analysis. However, the final analysis of age-specific fecundity was done excluding the last few age points (post-day40) to avoid an unbalanced design arising due to the death of most individuals towards the later age points. Additionally, the cumulative fecundity of females was also quantified across all the above-mentioned age points till death.

### Data analysis

The entire study used a total of eight populations. Populations with replicate identity (for example, BL_1_ and YLB_1_) were phylogenetically closer (see Figure S1) compared to those with different replicate identity. In addition, during an assay a given replicate set (for example, BL_1_ and YLB_1_) was handled together whereas different replicate sets were handled independently, often on different days. Thus data from replicate sets were treated as statistical blocks and was modelled as a random effect in all the analyses. The entire studythus involved four statistical blocks. Within each block, vials were the unit of replication and vial values / means were used as the unit of analysis / response variable. Except for adult survivorship and age-specific fecundity, results of all assays were analysed using mixed-model analysis of variance (ANOVA) on Statistica (version 10, Statsoft, Tulsa, OK, USA). In case of the analysis of juvenile traits, selection regime and sex (wherever applicable) were treated as fixed effects. For the early-life female fecundity assay and egg size measurement, selection regime and live-yeast concentration treatment were considered as fixed effects. For the analysis of the early-life progeny count data from Experiment2, male (selection) regime and female (selection) regime were used as fixed effects. For analysing female mean longevity data from Experiment3, selection regime and mating status were treated as fixed effects. Cumulative life-time fecundity data was analysed using selection regime as fixed factor.

For female survivorship analysis, we applied_a_ mixed Cox’s Proportional Hazard models, using the `coxme` package in R version 4.0.2 (Gupta et al. 2013, Therneau et al. 2015). The effects of regime, mating status and their interaction on survivorship was analysed and block was fitted as a random intercept. A few flies those escaped during the course of the assay were censored. Cox partial likelihood estimates were compared across treatment and selection regime.

Age-specific fecundity was analysed as linear mixed-effects models with the `lme4` package in R version 4.0.2 (Bates et al. 2014). Further, to obtain the p-values for the `lmer` models with degrees of freedom based on the Satterthwaite approximation, we used the R package `lmerTest` (Kuznetsova et al. 2017). Regime and time-point (age) were fitted as categorical fixed effects. Block as well as all interactions involving it were fitted as random effects. In addition, vial ID (replicate vial identity) was nested within block and was fitted as a random effect. The nested random effect accounts for the fact that subsequent egg counts from the same group of flies within a particular vial are not independent (Harrison et al. 2018). Post-hoc analysis was done using Tukey’s HSD with the `emmeans` package (Lenth 2018). To measure reproductive senescence, rates at which fecundity declined with age, after reaching a peak, were quantified for both regimes. Rate of decline in fecundity was quantified across seven time-points (*τ*), starting from the peak, i.e. day10 post-eclosion until day40. For a particular age-point, the mean of per-capita fecundity (*ε*) of a population was calculated using the observed fecundity across all the vials within that population. A linear model (*ε* ∼ b_p_*τ* + c_p_, where b_p_ is population specific slope and c_p_ is population specific intercept) was fitted to these data using the `lm` function in R to obtain a b-coefficient (i.e., slope) for each of the eight populations (Fig. S4 of supplementary information). The slope of the regression line served as a proxy for rate of reproductive senescence. Finally, these b-coefficients, without the sign, were analysed using mixed model ANOVA in Statistica, taking selection regime as a fixed effect and block as a random effect.

## 3. Results

### Experiment 1: Juvenile traits and female fecundity

We did not find a significant effect of selection regime on juvenile traits including development time, pre-adult (egg-to-adult) survivorship and dry body weight at the time of eclosion (Table S2). Analysis of the early-life fecundity data indicated significant effects of selection regime and yeast concentration (Table 1). The effect of selection regime × yeast concentration interaction was not significant (Table 1). Compared to the BL females, YLB females had 8.7-22.89 % higher mean per-capita fecundity across all live-yeast concentrations in the assay(Fig.1). Moreover, the higher early-life fecundity output of YLB females was consistent across a gradient of live-yeast concentration (Fig.1). The effect of selection regime on egg size was non-significant (Table S2). Hence, there was no evidence of differential provisioning in eggs produced by the experimental females.

**Table 1:** Results of the analyses on early-life fecundity(from experiment 1) and progeny output (from experiment 2) in females. Fecundity data was analysed using three factor mixed-model ANOVA with selection regime and live-yeast concentration as fixed factors, and block as random factor. Progeny count data from experiment 2 was analysed using three factor mixed-model ANOVA with female regime and male regime as fixed factors, and block as random factor. Statistically significant p-values are highlighted in boldface.

| Traits | Effects | SS | DF | MS | DEN<br>DF | DEN MS | F | p |
| --- | --- | --- | --- | --- | --- | --- | --- | --- |
| **Early life**<br>**fecundity** | Selection Regime (SR) | 947.8 | 1 | 947.8 | 3.01 | 7.47 | 126.83 | **<0.01** |
|  | Live-yeast concentration (YC) | 51192.3 | 2 | 25596.2 | 6.00 | 1043.57 | 24.52 | **<0.01** |
|  | Block | 19814.0 | 3 | 6604.7 | 5.59 | 1008.10 | 6.55 | **0.03** |
|  | SR × YC | 23.0 | 2 | 11.5 | 6.01 | 42.40 | 0.27 | 0.77 |
|  | SR × Block | 22.4 | 3 | 7.5 | 6.02 | 42.41 | 0.17 | 0.90 |
|  | YC × Block | 6266.1 | 6 | 1044.4 | 6.00 | 42.39 | 24.63 | **<0.01** |
|  | SR × YC × Block | 254.3 | 6 | 42.4 | 247.00 | 59.14 | 0.71 | 0.64 |
| **Progeny**<br>**count** | Female regime | 411.13 | 1 | 411.14 | 3.00 | 7.40 | 55.55 | **<0.01** |
|  | Male regime | 0.99 | 1 | 0.99 | 3.00 | 36.84 | 0.03 | 0.88 |
|  | Block | 3149.10 | 3 | 1049.70 | 0.64 | 20.48 | 51.26 | 0.26 |
|  | Female regime × Male regime | 7.18 | 1 | 7.18 | 3.00 | 23.76 | 0.30 | 0.62 |
|  | Female regime × Block | 22.20 | 3 | 7.40 | 3 | 23.76 | 0.31 | 0.82 |
|  | Male regime × Block | 110.51 | 3 | 36.84 | 3 | 23.76 | 1.55 | 0.36 |
|  | Fem. reg. × Male reg. × Block | 71.28 | 3 | 23.76 | 222 | 20.71 | 1.15 | 0.33 |

**Figure 1:**
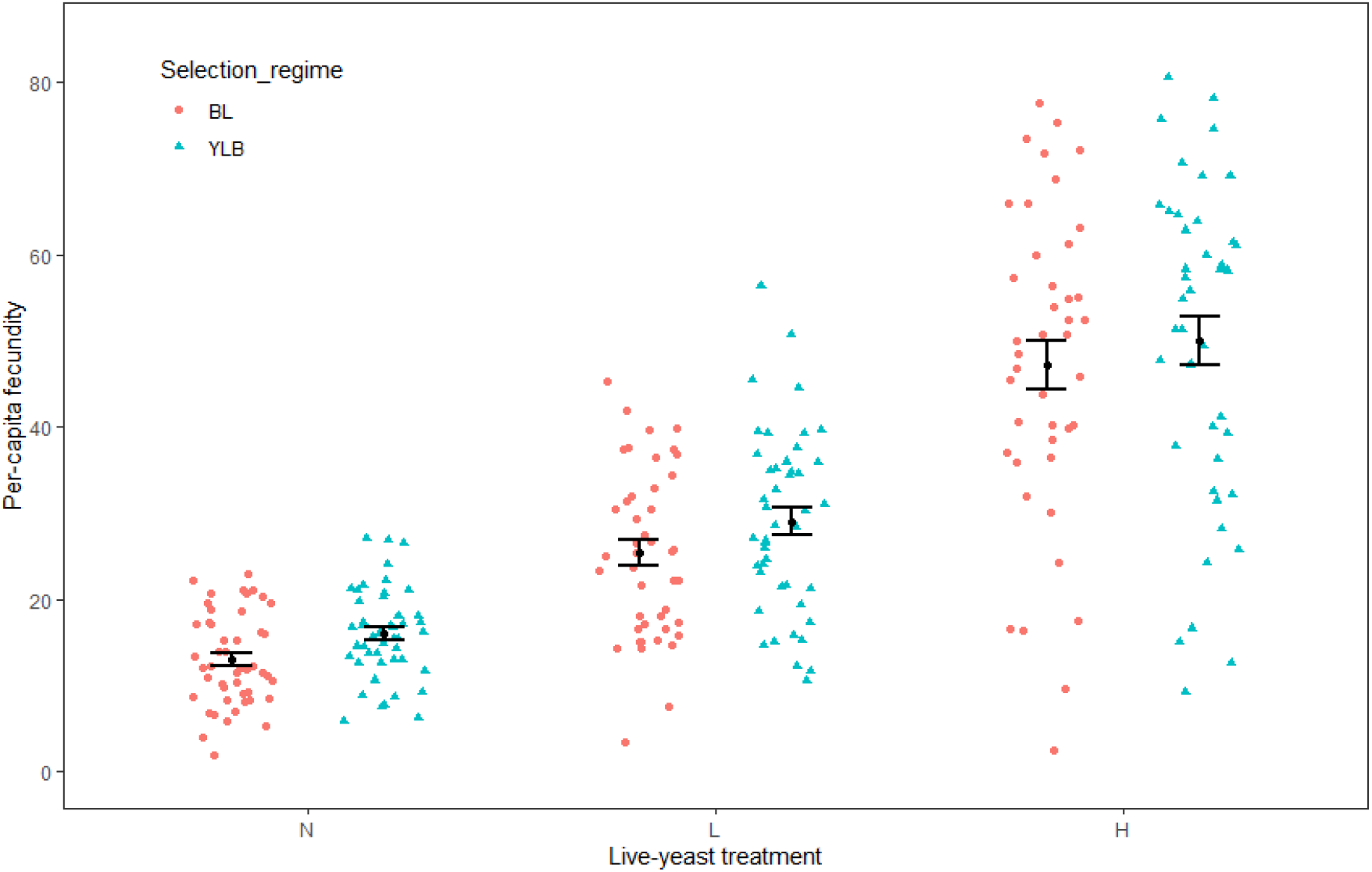
Female early-life fecundity from Experiment 1. Number of eggs laid during a 16-hour assay window by 4-5 day old (post-eclosion) females following the 48 hour live-yeast treatment was measured. This was done under three live-yeast conditions -no yeast (N), limiting yeast (L) and high yeast (H). The data distributions are shown as scatters. Pink circles represent data from BL (control regime) females, and teal triangles represent that from YLB (experimental regime). The black circle in each scatter represent the mean, which is shown along with the standard error (vertical error bars). Six of the twelve test females in a vial were used for fecundity measurement. Mean per-capita fecundity in a replicate vial was calculated using these six females. Three factor mixed model ANOVA indicated significant effects of selection regime and live-yeast treatment. However, the effect of selection regime × live-yeast treatment interaction was not significant.

### Experiment 2: Female × male effect on female fitness

The analysis of the early-life progeny output data from Experiment 2 indicated a significant effect female selection regime. Neither the male selection regime, nor the interaction between these two fixed factors were found to have significant effects on progeny count (Table 1, Fig. 2). YLB females produced significantly more progeny irrespective of the regime of the males they interacted with (Fig. 2). Thus, the higher early-life reproductive output in YLB’s seems to be independent of any male influence.

**Figure 2:**
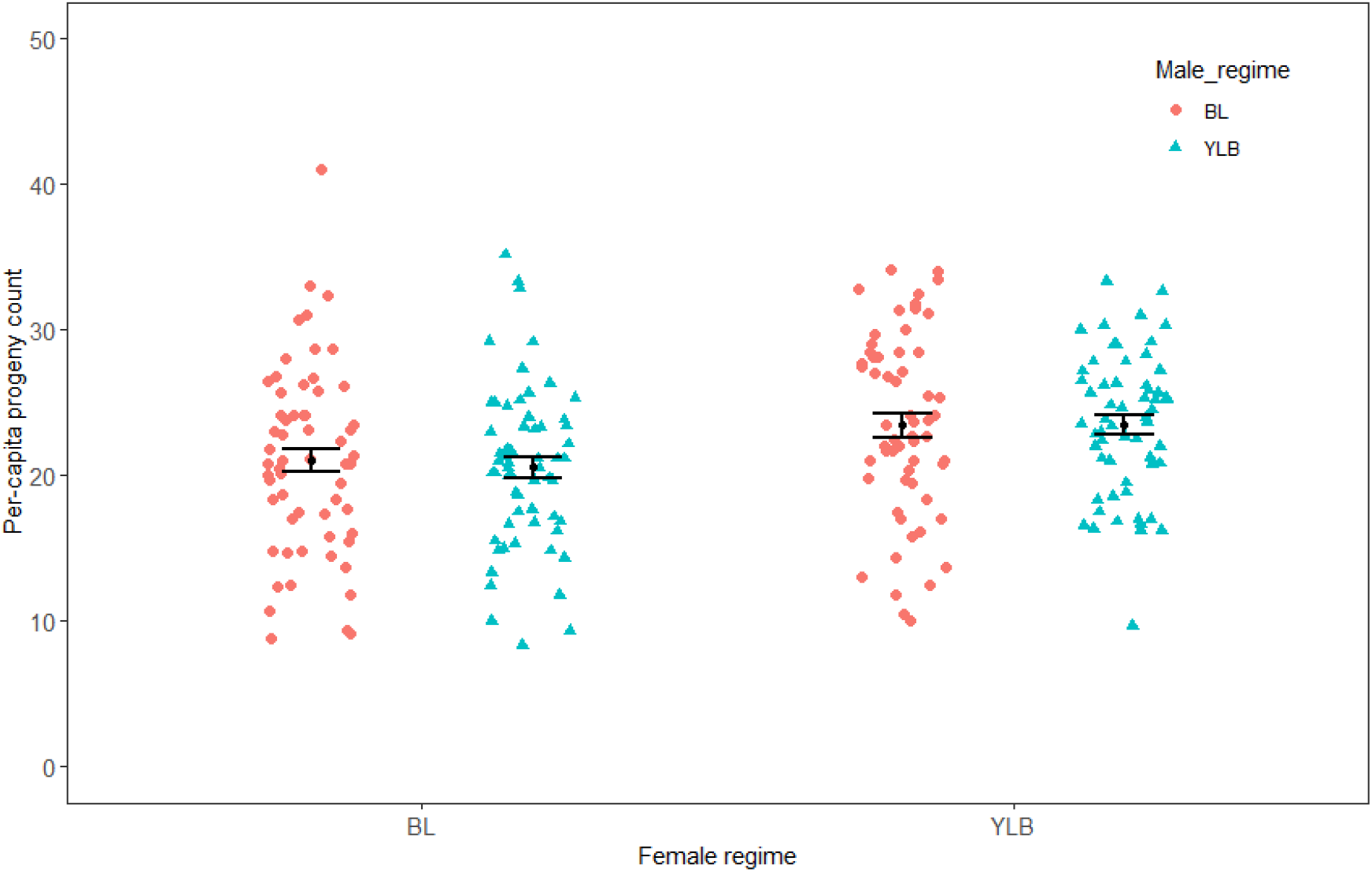
Female × male effect on early-life reproductive output in females as measured in Experiment 2. Number of progeny produced during a 17-hour assay window by 4-5 day old (post-eclosion) BL and YLB females, mated and held with BL and YLB males (full factorial combination), was measured. The data distributions are shown as scatters. Pink circles represent data from BL (control regime) females, and teal triangles represent that from YLB (experimental regime). The black circle in each scatter represent the mean, which is shown along with the standard error (vertical error bars). All six females in each replicate vial were allowed to produce progeny. Vials mean, which was calculated over these six test females, was used as the unit of analysis. Two factor mixed model ANOVA indicated a significant effect of female regime, but effect of male regime on progeny count was not significant, nor was that of the male regime × female regime interaction.

### Experiment 3: Longevity and age-specific fecundity

We did not find a significant effect of selection regime on survival probability (Table S4) and mean longevity (Table S5) of females. Mating status, however, had a significant impact. Virgin females had a better survival rate compared to the mated females (Fig. S3). Unsurprisingly, analysis of the mean longevity indicated that mating status had a significant effect on mean longevity (Table S5). Virgin females were found to have 67.58 % longer mean longevity compared to the mated females for both BL and YLB selection regimes (Fig. S4, Table S4). Results of these analyses can be found in the supplementary information.

Analysis of the cumulative fecundity of females quantified across different age points till death revealed that there was no significant effect of selection regime on per-capita fecundity, however, the effect of the selection regime×time point interaction was found to be significant (Table 3a, Fig.3a). Tukey’s adjusted p-value obtained by pairwise comparisons suggested that, as compared to the ancestors, YLBs have significantly higher fecundity at the age of day5 and day11 post-eclosion. Interestingly, there was a clear trend of YLBs having higher mean per-capita fecundity in initial age points compared to BLs (through age-5, 10, 15, 20 and 25 days post-eclosion). The trend, however, was found to be reversed at the later time points, with BLs showing comparatively higher fecundity (through age-30, 35 and 40 days post-eclosion). The detailed outcome of the post-hoc analysis is mentioned in the supplementary information (Supplementary sheet, Table-S6). Further, a comparison of the slopes (b-coefficients) of the linear regression fit of age-specific fecundity on time for the two regimes suggested a faster rate of decline in fecundity in YLBs compared to that of the BLs (mean ±SE: b_BL_=-2.03 ±0.41; b_YLB_=-3.36 ±0.13, Fig. S5). Two factor mixed model ANOVA on the b-coefficient indicated a significant effect of selection regime (Table 3b). Considering the b-coefficient to be a measure of reproductive senescence, YLB females showed faster reproductive senescence compared to that of the BL females. The cumulative fecundity of females, quantified across all the above-mentioned age points, did not differ between the two regimes (Fig. 3b, Table 3c)

**Table 3:** Results of the analyses of age-specific fecundity measured in Experiment 2. (a) Age-specific fecundity count was analysed using linear mixed-effect model, where selection regimeand time point (age) were fitted as categorical fixed effects, and block as random effect.(b) For each population, age-specific fecundity data was fitted in a linear model (*ε* ∼ b_p_*τ* + c_p_, where b_p_ is population specific slope and c_p_ is population specific intercept) to compute b-coefficient (slope). These b-coefficient values (i.e., slope values without the sign) were analysed using two factor mixed model ANOVA with selection regime and block as fixed and random factors respectively. (c) Cumulative fecundity, i.e., fecundity count summed over all age points, were analysed using a two factor mixed model ANOVA with selection regime and block as fixed and random factors respectively. Statistically significant p-values are mentioned in bold face.

| Trait | Effects | SS | DF | MS | Denominator<br>DF | F | p |
| --- | --- | --- | --- | --- | --- | --- | --- |
| (a) Age-specific fecundity | Selection Regime (SR) | 8.2 | 1 | 8.18 | 59.03 | 0.39 | 0.53 |
|  | Time point | 6531.9 | 7 | 933.13 | 22.27 | 44.44 | < **0.01** |
|  | SR × Time point | 986.2 | 7 | 140.88 | 22.41 | 6.71 | < **0.01** |
| (b) b-coefficient | Selection Regime | 4.1 | 1 | 4.1 | 3 | 13.68 | **0.03** |
|  | Block | 1.4 | 3 | 0.47 | 3 | 1.54 | 0.37 |
| (c) Cumulative Fecundity | Selection Regime (SR) | 35 | 1 | 35 | 3.01 | 0.06 | 0.82 |
|  | Block | 4089 | 3 | 1363 | 3 | 2.26 | 0.26 |
|  | SR × Block | 1806 | 3 | 602 | 76 | 0.73 | 0.53 |

**Figure 3:**
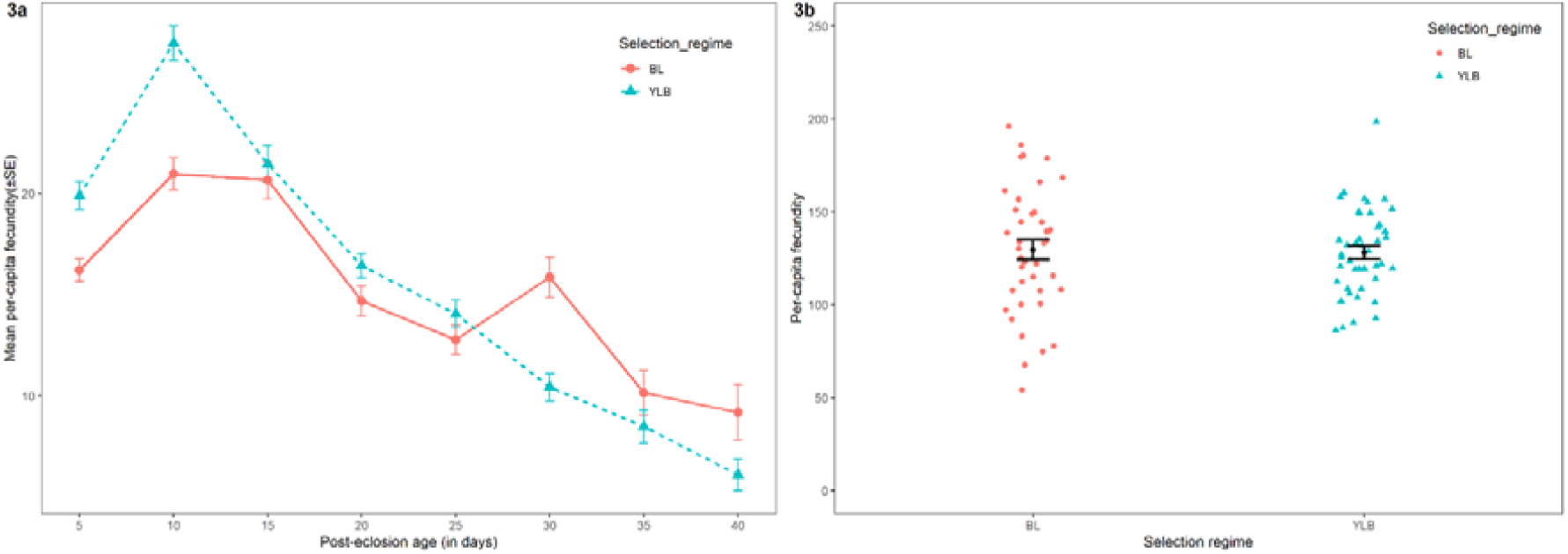
Age-specific and cumulative fecundity of females as measured in Experiment 3. Number of eggs laid during 17 hour windows on day 5, 10, 15, 20, 25, 30, 35, 40 were measured during the longevity assay. Only CE (continuously exposed) vials were used for this purpose. Per capita fecundity was calculated for a vial by diving the fecundity count by the number of surviving females, which was twelve to start with. (a) Mean age-specific per capita fecundity analysis indicated a significant effect of selection regime × age interaction. Pink circles represent data from BL (control regime) females, and teal triangles represent that from YLB (experimental regime). Data points represent mean per capita fecundity at the corresponding age point. The vertical error bars represent standard error of means. (b) The mean cumulative per-capita fecundity of females, quantified across all the above-mentioned age points. The data distributions are shown as scatters. Pink circles represent data from BL (control regime) females, and teal triangles represent that from YLB (experimental regime). The black circle in each scatter represent the mean, which is shown along with the standard error (vertical error bars). Analysis using two factor mixed model ANOVA indicated no significant difference in mean cumulative per-capita fecundity between the females of the two regimes.

## 4. Discussion

Yeast being the main source of protein, especially in a laboratory ecology (Chippindale et al. 1993, Lee et al. 2008), a culture protocol devoid of yeast supplement poses a significant deprivation of dietary protein, limiting reproductive output. Following twenty three generations of laboratory evolution under such a condition, we found clear evidence of evolution of fecundity traits indicating adaptive changes. Fecundity and progeny output in females most relevant to fitness showed a significant increase. However, there was no evidence to suggest a change in either lifetime reproductive out or lifespan. Instead, scheduling of reproductive output showed an interesting shift, wherein early life reproductive output was found to be significantly higher, with a faster drop as the females grew older, indicating faster reproductive senescence. Further, higher early life fecundity of females from the experimental regime was observed in absence of yeast supplement as well as in presence of low and high concentrations of the same, suggesting that plasticity in fecundity in response to variation in protein availability did not evolve. There are a number of possible ways in which females in the experimental populations could have attained higher early life reproductive output. We discuss these possibilities and the broader implication of our finding in the following section.

There are at least four different ways in which early-life fecundity and reproductive output could have increased in females. First, increased larval resource acquisition and/or storage can significantly contribute to the total resource available for reproduction, particularly early in life. In *D. melanogaster*, yeast-starvation at the larval stage dramatically brings down female fecundity such that females grown under complete yeast starvation can only remain reproductive for about 5-6 days, clearly indicating the importance of larval resource acquisition on early life reproductive output (Robertson & Sang 1944). However, we did not find any evidence upholding this theory. Neither juvenile traits (development time, egg-to-adult survivorship) nor body mass at eclosion was affected by experimental evolution. Alternatively, an increase in the relative abundance of different macronutrients, especially total protein content in adult females without affecting total body mass, could translate into increased fecundity. This idea of more fine scale physiological adaptation, however, remains to be tested with additional experimental data.

Secondly, number of offspring produced can be increased if investment in each offspring is reduced (Hein et al. 2018, also see Vijendravarma et al. 2010). We, however, did not find any evidence supporting such number vs. size trade-off as there was no effect of selection on egg size even though number of eggs produced by the females increased substantially. This is perhaps not surprising as previous investigations on *D. melanogaster* found egg size and number to be genetically uncorrelated (Schwarzkopf et al. 1999). Hence, egg production rate can evolve without affecting egg size.

A third possibility involves resource reallocation across life history traits such that higher amount of resources are made available to egg production. For example, resources invested in somatic maintenance could be reallocated to reproduction, especially when there is strong selection on higher reproductive output and little or no selection on longevity traits, such as in short lived organisms. Such a trade-off between reproduction and lifespan is pervasive in nature and has been widely reported even in laboratory systems (Rose 1984, Chippindale et al. 1993, Zwaan et al. 1995, Flatt 2011, Hansen et al. 2013, Maklakov et al. 2017). This theory is also inadequate in explaining our findings on fecundity evolution as we did not find a significant effect of experimental evolution on female longevity. This, however, is not surprising as several attempts to test reproduction-longevity trade-off have been either inconclusive or have failed (Le Bourg and Minois 1996, Dudash & Fenster 1997, Kotiaho & Simmons 2003, Partridge et al. 2005, Grandison et al. 2009, Zajitschek et al. 2016, 2019). There is a growing consensus that longevity and fecundity traits are, at least to some extent, genetically independent of each other and the trade-off between them can be uncoupled (reviewed in Flatt 2011). Importantly, in the present investigation we did not find any evidence suggesting change in total lifetime reproductive output. If reproduction-longevity trade-off is more sensitive to the total lifetime reproductive effort, a lack of response in the longevity traits is not unconceivable.

A fourth, and perhaps the most plausible scenario is the change in reproductive schedule without any alteration in life time reproductive output. Importantly, due to the 14 day discrete generation cycle, early life fecundity is a major determinant of female fitness in our laboratory ecology (see similar arguments in Promislow and Tatar 1998, Houle and Rowe 2003, Prasad & Joshi 2003, Rice et al. 2006). Both survival and reproduction beyond day14 of the generation cycle (i.e., approximately 4-5 days of adult age) have no contribution to fitness. Hence, early life fitness components, including fecundity, are exclusive targets of selection under such an ecology. This view of the population demography would predict the evolution of early life fitness regardless of its consequences to the late life performance (Stearns 1992). This theory fits perfectly with our age specific fecundity results, wherein females from the evolved populations were found to have significantly higher fecundity. The associated observation of a faster decrease in fecundity without a significant change in mean lifetime egg output seems to fit antagonistic pleiotropy theory of ageing (Williams 1957). Early investigations on *Drosophila* populations substantiated the theory (Rose 1984, Rauser et al. 2006), and have since been shown to explain both somatic as well as reproductive senescence. Multiple investigations also reported a substantial genetic variation in age-specific fecundity in *D. melanogaster* (for example, Tatar et al. 1996, Durham et al. 2014). More recently, using *C. elegans*, Anderson et al. (2011) found that genetically heterogeneous populations selected for early-life fecundity evolved increased early but reduced late-life reproductive output without suffering a total fitness or lifespan cost. Thus, it is not unreasonable to suggest that adaptation seen in our YLB populations involves the evolution of early-life beneficial alleles with detrimental pleiotropic effects on late-life reproductive output.

Adaptive evolution of early-life fecundity in YLB females seems to support the hitherto reported phenomenon of the release of hidden genetic variation in novel ecology or stress (Badyaev 2005). Several past investigations have shown that novel environmental challenges or stresses (e.g., heat shock) can generate new trait variances or convert otherwise phenotypically neutral alleles to non-neutral (Hall 1990, Ratner et al. 1992, Kondrashov and Houle 1996, Wessler 1996, Leips & Mackay 2000, Queitsch et al. 2002, Dworkin et al. 2003, Nedelcu & Michod 2003, Ruden et al. 2003, Sollars et al. 2003). Such novel genetic variance can then be subjected to selective pressures leading to adaptation. Further, there is a growing literature on substantial G×E interaction on fitness-related traits, including fecundity (see Leroi et al. 1994, Fry et al. 1998, Burns et al. 2012, Pallares et al. 2021). Most notably, Pallares et al. (2021) reported that dietary sugar enrichment to have a profound effect on the fitness landscape of the fecundity and longevity traits in *D. melanogaster*. Fitness consequences of many alleles were found to be much stronger in the altered diet compared to the standard diet (Pallares et al. 2021). Our results could also be affected by a similar diet dependence of fitness consequence of fecundity traits in this system. Alleles that affect unstimulated (by live yeast supplement) fecundity could be selectively neutral in the ancestral BL populations, but have strong fitness effects under yeast deprived ecology of the YLB populations and hence, be subjected to directional selection.

The identity of physiological traits that could be subjected to selection during such an adaptation would depend on the proximate processes affecting early life fecundity. Therefore, the proximate mechanistic explanation of higher early-life fecundity in the evolved females is an important issue. We set aside this problem as dealing with it will require additional data. However, there are several plausible theories to explore. For example, increased foraging behaviour (McCaffery 1975, Ives 1981, Barnes et al. 2008), reduction in wastage of resources in non-essential activities such as, spontaneous locomotion (Long & Rice 2006), better resource recycling (Scott et al. 2004, Barth et al. 2011, Cuervo & Macian 2012, Adler & Bonduriansky 2014, Mason et al. 2018) can all lead to increased resource availability for egg production in females.

As discussed earlier, in case of short term food shortage, the strategy to divert resources away from reproduction so that the temporary famine can be overcome, as suggested by adaptive resource reallocation theory, could be adaptive. Here we have shown that selection can favour novel reproductive strategies that might involve increased reproduction, if the famine is long lasting. For a short lived species, as our results suggests, that could lead to increased reproductive output at the cost of late life fertility.

## 6. Acknowledgements

We thank Daniel E. L. Promislow and Benjamin Harrison of University of Washington for their constructive criticism and suggestions on an earlier version of the manuscript. Integrating their suggestions have greatly improved the quality of the manuscript, including the clarity of the presentations. We thank the two anonymous reviewers of a previous version of the manuscript for their immensely helpful and constructive criticisms. Incorporation of their suggestions have greatly improved the quality of the manuscript. We are thankful to Tanya Verma for her help during the experiments and population maintenance. The experiments reported here was supported by the Core Research Grant (file no.

CRG/2019/002460) from Science and Engineering Research Board, Department of Science and Technology, Government of India. PD thanks Council for Scientific and Industrial Research, Government of India for financial support in the form of Junior and Senior Research Fellowship. NP thanks Summer Research Fellowship Programme of Jawaharlal Nehru Centre for Advanced Scientific Research, Bengaluru for financial assistance during his summer internship.

## Notes

### Competing Interest Statement

The authors have declared no competing interest.

### Summary of Updates

This is a heavily revised version. Introduction and Discussion sections have been heavily rewritten. There are a few new analysis and figures in the manuscript and the supplementary information file. The title has been modified to represent the most important finding understandable to a broader audience. There is no additional data in the revised version.

