## Supplementary material for "Evolution of a novel female reproductive strategy in *Drosophila melanogaster* populations subjected to long term protein restriction": Supplentary information file

**Supplementary Information**

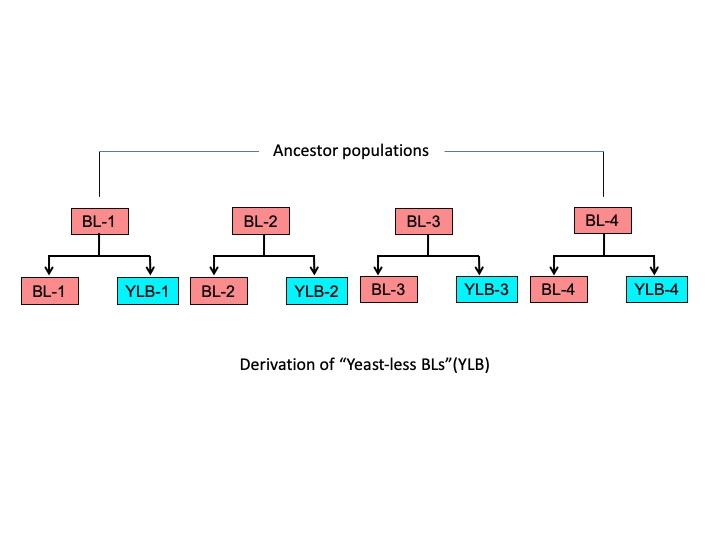

Figure S1: A schematic representation describing the derivation and the ancestry of the ‘Yeast-less BLs’ (YLB)

| **Mix components** | **Ingredients** | **Amount** |
| --- | --- | --- |
| Food paste | a. Banana | 205 g |
|  | b. Jaggery | 35 g |
|  | c. Barley | 25 g |
|  | d. Baker’s yeast | 36 g |
|  | e. Water (to prepare food paste) | 180 mL |
|  | f. Absolute Ethanol (to mix with Yeast) | 22 mL |
| Agar mix | g. Water (to mix with Agar-agar) | 1 L |
|  | h. Agar-agar | 12.4 g |
| Preservative | i. p-Hydroxy methyl benzoate | 2.4 g |
|  | j. Absolute Ethanol (to dissolve benzoate) | 23 mL |

Table S1: Ingredients for the preparation of one litre of standard banana-sugarcane jaggery- yeast medium. Banana, jaggery, and barley are grounded together with 180 mL of water. Live yeast and absolute ethanol are added and mixed until the granules dissipate. The whole mixture is then added to the boiling agar-agar in 1L of water. The food is boiled properly. p-hydroxy methyl benzoate is mixed with absolute ethanol and added to the medium once the temperature falls below 60°C.

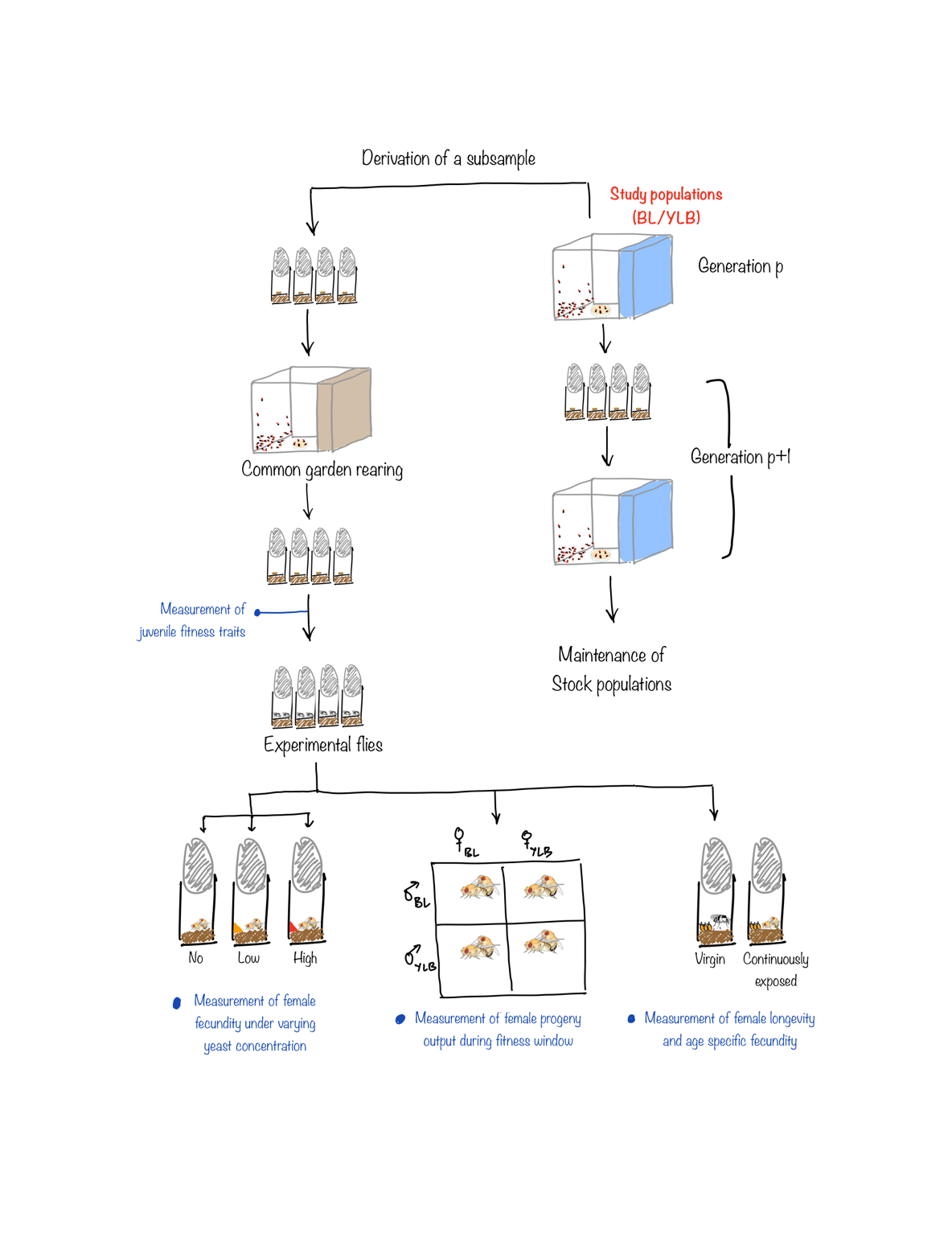

Figure S2: Schematic representation of the general plan of the study. All assays were performed on experimental flies raised from a subset of experimental populations. Raising of the experimental flies included one generation of common garden rearing that matched the ecology of the ancestral rearing regime. Three separate assays (Experiment 1, 2 and 3 described in the text) were performed using the experimental flies.

| **Traits** | **Effects** | **SS** | **DF** | **MS** | **DEN DF** | **DEN MS** | **F** | **p** |
| --- | --- | --- | --- | --- | --- | --- | --- | --- |
| **Pre-adult Development time** | Selection Regime (SR) | 2 | 1 | 2 | 3.01 | 6.05 | 0.36 | 0.59 |
|  | Sex | 856 | 1 | 856 | 3.07 | 0.92 | 929.32 | **<0.01** |
|  | Block | 1555 | 3 | 518 | 2.72 | 5.91 | 87.71 | **<0.01** |
|  | SR × Sex | 5 | 1 | 5 | 3.06 | 1.06 | 4.47 | 0.12 |
|  | SR × Block | 18 | 3 | 6 | 3 | 1.05 | 5.77 | 0.09 |
|  | Sex × Block | 3 | 3 | 1 | 3 | 1.05 | 0.87 | 0.54 |
|  | SR × Sex ×Block | 3 | 3 | 1 | 138 | 6.40 | 0.16 | 0.92 |
| **Juvenile survivorship** | Selection Regime (SR) | 0.001 | 1 | 0.001 | 3 | 0.01 | 0.09 | 0.78 |
|  | Block | 0.004 | 3 | 0.001 | 3 | 0.01 | 0.12 | 0.94 |
|  | SR × Block | 0.04 | 3 | 0.01 | 69 | 0.005 | 2.29 | 0.09 |
| **Dry (body) weight at eclosion** | Selection Regime (SR) | 1.43×10^-6^ | 1 | 1.43×10^-6^ | 3.00 | 4.63×10^-4^ | 3.08×10^-3^ | 0.96 |
|  | Sex | 0.27 | 1 | 0.27 | 3.00 | 9.11×10^-4^ | 294.33 | **<0.01** |
|  | Block | 7.43×10^-3^ | 3 | 2.48×10^-3^ | 4.57 | 1.27×10^-3^ | 1.95 | 0.25 |
|  | SR × Sex | 1.71×10^-5^ | 1 | 1.71×10 ^-5^ | 3.01 | 1.06×10^-4^ | 0.16 | 0.71 |
|  | SR × Block | 1.39×10^-3^ | 3 | 4.63×10^-4^ | 3 | 1.06×10^-4^ | 4.39 | 0.13 |
|  | Sex × Block | 2.73×10^-3^ | 3 | 9.11×10^-4^ | 3 | 1.06×10^-4^ | 8.63 | 0.05 |
|  | SR × Sex × Block | 3.17×10^-4^ | 3 | 1.06×10^-4^ | 141 | 3.46×10^-4^ | 0.30 | 0.82 |
| **Egg size** | Selection Regime (SR) | 6.18 × 10^4^ | 1 | 6.18 × 10^4^ | 3.0060 | 3.27 × 10^6^ | 0.02 | 0.89 |
|  | Live-yeast concentration (YC) | 9.98 × 10^7^ | 2 | 4.98 × 10^7^ | 6.0139 | 4.03 × 10^6^ | 12.38 | **0.007** |
|  | Block | 4.63 × 10^7^ | 3 | 1.54 × 10^7^ | 5.0000 | 4.05 × 10^6^ | 2.49 | 0.11 |
|  | SR × YC | 8.87 × 10^6^ | 2 | 4.43 × 10^6^ | 6.0049 | 1.14 × 10^7^ | 0.39 | 0.69 |
|  | SR × Block | 9.81 × 10^6^ | 3 | 3.27 × 10^6^ | 6.0022 | 1.14 × 10^7^ | 0.29 | 0.83 |
|  | YC × Block | 2.41 × 10^7^ | 6 | 4.03 × 10^6^ | 6.0000 | 1.14 × 10^7^ | 0.35 | 0.88 |
|  | SR × YC × Block | 6.85 × 10^7^ | 6 | 1.14 × 10^7^ | 256.000 | 6.99 × 10^6^ | 1.63 | 0.14 |

Table S2: Results of the analyses on traits measured in Experiment 1 (development time, pre-adult survivorship, dry body weight at eclosion and egg size). Egg-to-adult development time and dry body weight at eclosion data were analysed using three factor mixed-model ANOVA with selection regime and sex as fixed factors, and block as random factor. Pre-adult survivorship (i.e., egg-to-adult survival rate) was analysed using two factor mixed-model ANOVA with selection regime as fixed factor, and block as random factor. Egg size data was analysed using three factor mixed-model ANOVA with selection regime and live-yeast concentration treatment as fixed factors, and block as random factor. Statistically significant p-values are highlighted in boldface.

| **Traits** | **Selection regime** | **Sex** | **Yeast conc.** | **Mean** | **Std. Err.** |
| --- | --- | --- | --- | --- | --- |
| **Pre-adult development time**  **(in hours)** | YLB | Female | - | 216.83 | 0.40 |
|  | YLB | Male | - | 221.20 | 0.40 |
|  | BL | Female | - | 216.24 | 0.41 |
|  | BL | Male | - | 221.32 | 0.41 |
| **Juvenile survivorship** | YLB | - | - | 0.74 | 0.01 |
|  | BL | - | - | 0.73 | 0.01 |
| **Dry weight at eclosion**  **(in mg)** | YLB | Female | - | 0.338504 | 0.002982 |
|  | YLB | Male | - | 0.255120 | 0.002982 |
|  | BL | Female | - | 0.338033 | 0.002941 |
|  | BL | Male | - | 0.255972 | 0.002982 |
| **Egg size**  **(μm^2^)** | YLB | - | N | 73514.41 | 385.81 |
|  | YLB | - | M | 72526.18 | 385.81 |
|  | YLB | - | H | 72596.12 | 390.07 |
|  | BL | - | N | 73913.82 | 395.13 |
|  | BL | - | M | 72076.24 | 381.49 |
|  | BL | - | H | 72735.87 | 385.81 |

Table S3: Descriptive statistics of traits measured in Experiment 1. Development time, pre-adult survivorship, dry body weight at the time of eclosion and egg size are mentioned here. Values of the rest of the traits can be found in the main text.

| **Fixed coefficients** | **Hazard ratios** | **Lower CI** | **Upper CI** | **z** | **p** |
| --- | --- | --- | --- | --- | --- |
| Selection Regime: YLB | 0.99 | 0.88 | 1.12 | -0.08 | 0.94 |
| Mating status: Virgin | 0.13 | **0.11** | **0.15** | -29.57 | **<0.01** |
| Sel. Reg. YLB × Mating status Virgin | 0.95 | 0.80 | 1.13 | -0.58 | 0.56 |
| **Random effects** | **Variance** |  |  |  |  |
| Block | 0.019 |  |  |  |  |

Table S4: Results of the cox proportional hazard analysis on female survivorship under virgin and mated conditions. Hazard rates are expressed relative to the hazard rate of default level of each fixed factor constrained to be 1. The default level of selection regime is ‘BL’ while that of mating status is ‘mated’. Lower CI and Upper CI indicate lower and upper bounds of 95% confidence intervals. Confidence intervals that do not contain 1 signify statistical significance.

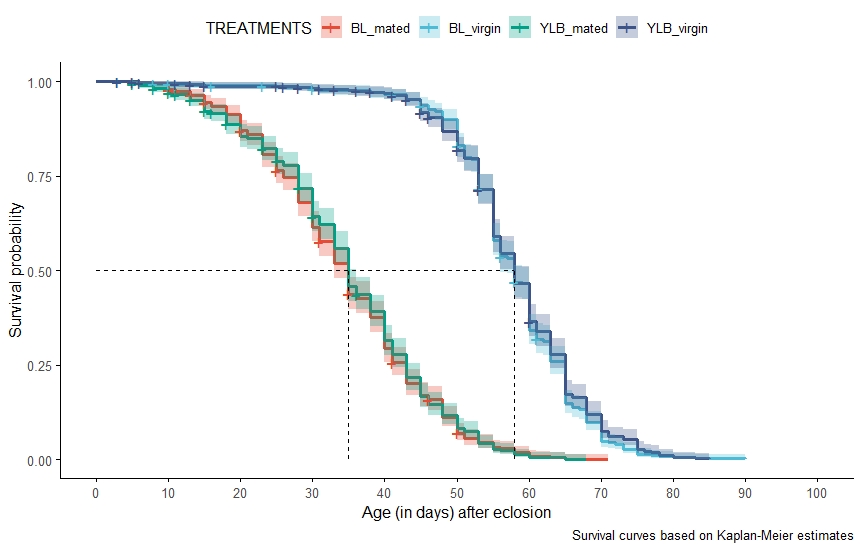

Fig. S3: Survival plots from Experiment 3. Survival curves were plotted using Kaplan – Meier Method. The difference between ‘mated’ and ‘virgin’ condition was found to be statistically significant. However, within each mating status (i.e., mated and virgin), the effect of selection regime was not significant.

| **Effects** | **SS** | **DF** | **MS** | **DEN DF** | **DEN MS** | **F** | **p** |
| --- | --- | --- | --- | --- | --- | --- | --- |
| Selection Regime (SR) | 15.17 | 1 | 15.17 | 3.00 | 18.70 | 0.81 | 0.43 |
| Mating Status (MStat) | 23067.32 | 1 | 23067.32 | 3.00 | 48.47 | 475.86 | **<0.01** |
| Block | 123.43 | 3 | 41.14 | 2.03 | 46.11 | 0.89 | 0.56 |
| SR × MStat | 0.05 | 1 | 0.05 | 3.00 | 21.08 | 0.002 | 0.96 |
| MStat × Block | 145.47 | 3 | 48.49 | 3 | 21.08 | 2.30 | 0.25 |
| SR × Block | 56.11 | 3 | 18.70 | 3 | 21.08 | 0.88 | 0.54 |
| MStat × SR × Block | 63.24 | 3 | 21.08 | 155 | 19.47 | 1.08 | 0.36 |

Table S5: **Result of the analysis of mean longevity of females measured in Experiment 3**. Mean longevity was analysed using a three factor mixed model ANOVA with selection regime and mating status as fixed factors, and block as random factor. Statistically significant p-values are mentioned in bold face.

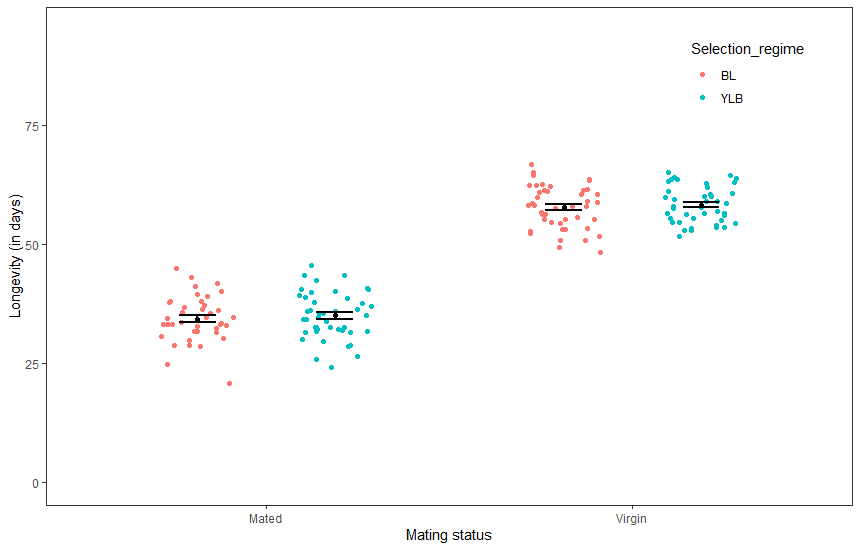

Fig. S4: Mean longevity of the females as measured in Experiment 3. Mean longevity was calculated for a replicate vial having twelve females. These values were then used as the unit of analysis. The data points in the plot represents these values. The data points have been jittered to increase visibility. Mean and standard error are shown. Analysis using three factor mixed model ANOVA indicated a significant effect of mating status, but the effect of selection regime and that of selection regime × mating status interaction was not significant.

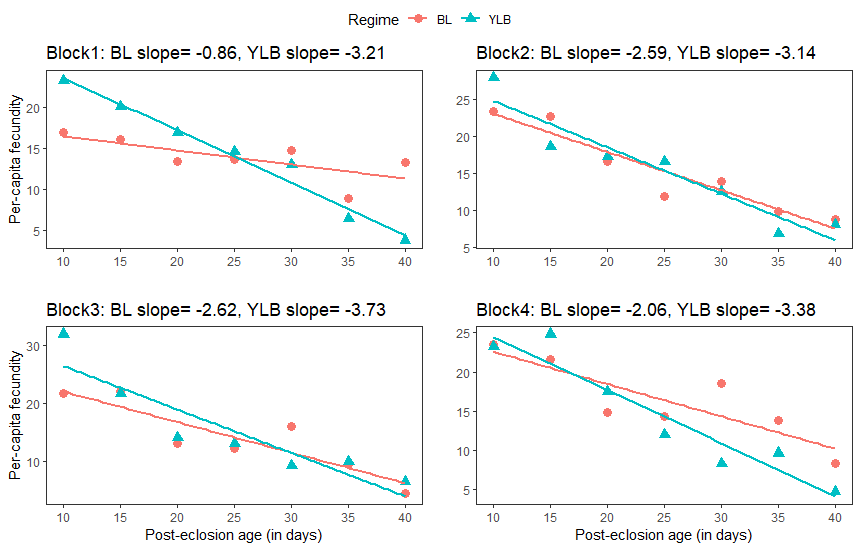

Fig S5: **Linear regression fit of age-specific fecundity.** Slope(b-coefficient) of the linear decline in per-capita fecundity with age following the peak fecundity was computed for each population by fitting a linear model to the mean age-specific per-capita fecundity. The two factor mixed model ANOVA on the b-coefficients indicated a significant effect of selection regime.

| **Age points** | **p value** |
| --- | --- |
| D-05 | **0.02** |
| D-10 | **<0.01** |
| D-15 | 0.7 |
| D-20 | 0.24 |
| D-25 | 0.39 |
| D-30 | **<0.01** |
| D-35 | 0.27 |
| D-40 | 0.07 |

Table S6**:** Outcome of the post-hoc analysis of age-specific fecundity comparison between BL and YLB. Tukey’s adjusted p-values obtained by pairwise comparisons suggested that, as compared to the ancestors, YLBs have significantly higher fecundity at the age of day 6 and day 11 post eclosion which drops significantly to a lower value at day 31. Statistically significant p-values are mentioned in bold face.

| **Age-specific fecundity** | PCF ~ REGIME+ TIME+ REGIME:TIME + (1\|BLOCK/VIAL_ID) + (1\|BLOCK:REGIME) + (1\|BLOCK:TIME) + (1\|BLOCK:TIME:REGIME) |
| --- | --- |
| **Female survivorship** | coxme (Surv (TIME, CENSOR)) ~ REGIME + MATING_STATUS +  REGIME : MATING_STATUS + (1\|BLOCK) |

Table S7:Final models used for the analyses on age-specific fecundity and female survivorship using R.
